# What’s encoded in a marmoset phee call? Food context beyond arousal and valence

**DOI:** 10.64898/2026.07.09.737477

**Authors:** Elodie F. Briefer, Kaja Wierucka, Florine A. Ermatinger, Rahel K. Brügger, Eva Ciccarelli, Kristin Meshinska, Karina Stampe Ernst, Judith M. Burkart

**Affiliations:** Behavioural Ecology Group, Section for Ecology and Evolution, University of Copenhagen, Copenhagen, Denmark; German Primate Center - Leibniz Institute for Primate Research, Behavioral Ecology and Sociobiology Unit, Kellnerweg 4, 37077 Göttingen, Germany; Department of Evolutionary Anthropology, University of Zurich, Winterthurerstrasse 190, 8057 Zurich, Switzerland; Institute of Agricultural Sciences, ETH Zürich, Universitätsstrasse 2, 8092 Zürich, Switzerland; Faculty of Health and Medical Sciences, University of Copenhagen, Blegdamsvej 3B, 2200 Copenhagen, Denmark; Center for the Interdisciplinary Study of Language Evolution (ISLE), University of Zurich, Affolternstrasse 56, 8050 Zürich, Switzerland

**Keywords:** bioacoustics, expression of emotions, machine learning, non-human primate

## Abstract

Animal vocalisations can convey information about external events, but whether this goes beyond reflecting the emotional state elicited by these events is debated. To explore this, we studied the acoustic structure of common marmoset (Callithrix jacchus) phee (long-distance contact) and ek (alert/mobbing) calls produced in five treatments varying in the emotional valence and arousal they elicit (internal state), as well as food and social context (external events). We measured changes in arousal via nasal temperature and analysed both basic acoustic parameters and Mel-frequency cepstral coefficients (MFCCs) of the calls. Support Vector Machines combined with Linear Mixed effect models revealed that phee calls encode both external events and internal states, while eks reflected predominantly arousal. Notably, an acoustic signature related to food context was present in phees both when provided (positive valence) and teased with highly preferred food items (negative valence), and even when food was not physically present (food call playback treatment). This suggests marmoset long-distant phee calls encode external information beyond emotional arousal and valence, and independently of the presence of an immediately triggering stimulus.

## Introduction

Non-human animal (hereafter ‘animal’) vocalisations were originally considered mere reflections of the current internal state of the caller, thus only conveying information about emotional state and other involuntary cues (e.g. physical attributes [1]). Accordingly, there is ample evidence now that vocalisations are tightly linked to the emotional state of the caller, in both human and non-human animals [2–5]. The later discovery of vocalisations that inform about events or objects in the external world (e.g. predators or food) revolutionised the field of animal communication [6]. Nevertheless, more than 40 years later, it remains unclear whether the proximate mechanisms underlying the production of calls or call parameters referring to external events are similar to human semantic communication, and hence constitute signs referring to objects or events independently of the caller’s emotional state. In particular, we are still missing compelling experimental evidence showing that the encoding of information about external events in vocalisations does not simply arise as a by-product, because the presence of the external event or stimulus triggers a specific emotional state in the caller that determines its vocalisations.

Among primates, common marmosets (Callithrix jacchus) stand out with considerable vocal flexibility and control [7–12]. Their extensive vocal repertoire encodes diverse information, including population, group membership, sex, and individual identity [13–20]. They produce not only simple tonal signals but also complex calls with multiple frequency components [21]. Moreover, they can form call combinations of up to nine distinct call types [14, 15, 22, 23], and show evidence for babbling and vocal production learning [11, 19, 24]. Importantly, common marmosets show vocal control with regard to when, where, and what to vocalise [e.g. reviewed in 25, 10, 26, 12]. For instance, they can change the duration [27], intensity ([27, 28], but see [29]), complexity [30], and the timing [29, 31] of calls. They are also capable of modulating calls even if the onset of environmental disruption occurs after call initiation [29]. In addition, their calls have been shown to inform about external events, such as the presence of food [32] or specific individuals [20]. However, it is unclear how much of this communication about external events could in fact reflect the internal state of the caller. Liao et al. [33] achieved a first step in answering this question by investigating the relationship between vocal output and arousal (assessed through heart rate) in marmosets. They found that the types and acoustic structure of marmoset vocalisations reflect more than arousal and are also linked to external events, such as the interindividual distance and timing of the partner’s vocalisation.

Here, we investigated whether common marmoset vocalisations encode information about external events, even when controlling for the variation linked to the emitter’s emotional states, and considering not only emotional arousal (bodily activation), but also emotional valence (i.e. whether the emotion is pleasant/positive or unpleasant/negative). We specifically focussed on the acoustic structure of call types that are known to be emitted across contexts rather than call type usage, as our aim was to differentiate referential from emotional content, the latter being typically expressed as variation within the acoustics structure of calls [3, 34]. Our analyses were thus carried out on marmoset ‘phee’ (long-distant contact) and ‘ek’ (alert/mobbing) calls, which can be emitted in several contexts [13]. We exposed marmosets to five situations (treatments) that triggered various levels of arousal (assessed through infrared thermography [35]), and differed in their emotional valence (negative, neutral, positive). In addition, the treatments differed depending on whether they were related to food or not, and whether they included a social component or not (presence vs. absence of conspecific call playbacks) (Table 1).

**Table 1.** Test treatments under which common marmoset vocalisations were recorded. Arousal varied—and was assessed—on an individual level based on changes in nasal temperature relative to baseline. Levels shown here correspond to averages across individuals. The arousal levels were based on significant temperature changes for each individual. They do not exactly match values shown in (**?**), as these averages are restricted to individuals that called during the experiments and could thus be included in our database.

| Treatment | Treatment description | Temperature change | Arousal level* | Valence | Food context | Social context |
| --- | --- | --- | --- | --- | --- | --- |
| PB— | Playback of agonistic calls | −0.71 | 0.38 | Negative | No | Yes |
| Teasing | Food teasing | −0.68 | 0.88 | Negative | Yes | No |
| Control | No stimulus | 0.00 | −0.22 | Neutral | No | No |
| PB+ | Playback of food calls | −0.28 | 0.50 | Positive | Yes | Yes |
| Food | Providing preferred food | 0.15 | −0.25 | Positive | Yes | No |

\* Corresponding arousal level based on average individual changes in nasal temperature relative to baseline.

Our predictions were as follows; first, if emotional arousal and valence (i.e. internal states) are the main driver of call acoustic variation between our treatments, we expected call parameters to change with arousal, and/or to differ between positive and negative contexts. More specifically, based on common patterns of vocal expression of emotions across species [2, 3], we expected calls to become higher in frequencies and have a more variable fundamental frequency with increasing arousal, while we expected calls produced in the positive treatments to be shorter and have lower and less variable fundamental frequencies than those produced in the negative treatments. Given that phees are long-distance contact calls that occur in both positive (e.g., reunion) and negative (e.g., separation/isolation, between-group encounters) contexts [14], we expected them to encode both arousal and valence. In contrast, since eks are typically produced in ambiguous-negative, mild to high-arousal situations (e.g. response to threats, mobbing/aggression), we expected them to primarily reflect arousal. In contrast to our first prediction, if external factors are the main driver of call acoustic variation between our contexts, we expected call structure to significantly differ between food vs. non-food related contexts, and/or between social (i.e. during conspecific call playbacks) vs. non-social contexts (i.e. in the absence of playbacks). We predicted that external factors would particularly influence the vocal structure of the highly flexible phee calls, which are also of longer duration than ek calls. Our two hypotheses are of course non-exclusive, as it is likely that animal calls simultaneously encode both emotional and referential information, similarly to human voice [36, 37]. In that case, marmoset call structure could vary simultaneously with internal states and external factors. Importantly, since we concurrently tested the effect of both types of factors, and since they varied relatively independently from each other across our contexts (Table 1), we were able to test if the information about external factors persists even after variation linked to internal states is accounted for. Such an outcome would suggest that common marmoset calls convey information about external factors beyond their current emotional state.

## Methods

The vocalisations used in this study were recorded as part of an experiment carried out and previously described in [35]. We thus only provide here a summary of the methods (for more details, please refer to [35]).

### Subjects and housing

The subjects were 17 adult common marmosets, nine females and eight males (Callithrix jacchus), that were born and raised in captivity. They were housed indoor in breeding pairs or family groups of up to seven individuals, with access to outdoor enclosures. They were aged 1 to 14 years at the time of the study.

### Experimental procedure

The subjects were tested within a testing compartment (60×50×50 cm) made of mesh situated within a testing room next to their home enclosure, which they had been habituated to over several weeks. The monkeys entered the testing compartment voluntarily via a tube system connecting their home enclosure to the testing room. The experimental session started when the subject was alone in the testing compartment and did not show any sign of moderate/high arousal. Each experimental session was divided into a baseline (hereafter “pre-test”), a stimulation (“test”) and a post-stimulation (“post-test”) phase. During the pre-test phase (2–3 min), the experimenter offered a small amount of food (mealworm or a small piece of nut cookie) to the subject every 20 s. This phase allowed us to obtain baseline values for the collected data (nasal temperature and behaviour [35], vocalisations in the current study). The test phase started 20 s after the last food item was handed over to the animal. It lasted 21–60 s (see [35] for the duration of each test phase for each subject) and consisted of one of the five treatments described in the section below. The post-test phase was identical to the pre-test phase and lasted about 3 min. During the test phase, the marmosets were submitted to the five test treatments listed in Table 1 and described above. They were each presented to the subjects on a different day and in a randomised order.

### Determination of the emotional valence and arousal

#### Valence

The emotional valence (positive vs negative) of the treatment was determined based on intuitive inference [39], following [35]. Negative emotions are triggered by contexts that would decrease fitness in natural life; the treatments Teasing and PB-, were thus assumed to be negative. By contrast, positive emotions occur in situations contributing to increased fitness in natural life; the treatments Food and PB+ were hence assumed to be positive for the animals. Finally, the control treatment was assumed to be relatively neutral.

#### Arousal

The level of arousal (bodily activation) of each subject during each treatment was assessed based on nasal temperature measurements [35]. In our previous study [35], and similarly to previous studies on other primates (e.g. chimpanzees [49]; macaques [50]; see review table in [35]), nasal temperature was validated as a good indicator of arousal; a decrease in temperature occurred when marmosets were exposed to negative arousing stimuli, independently of physical activity, and correlated negatively with piloerection of the tail, which was used as an independent arousal indicator [35]. To summarise the procedure, these measurements were extracted from video frames collected with an infrared thermography camera of the model FLIR T620 (temperature sensitivity: 0.04°C; resolution: 640×480 pixels; sampling rate: 30 fps). Using a customized MATLAB (R2018b) script, the minimal temperature of the region of interest (nose) on the first appropriate frame (defined based on 4 criteria, see in [35]) in each two-second interval was manually extracted. We chose minimal temperature instead of the average because the marmosets were able to freely move in the experimental cage. This made it impossible to position an ellipse to cover a region of interest of the same size in each frame and therefore causing an average to be inadequate. Then, to measure the temperature changes relative to baseline, we used baseline centred temperature values. To calculate these values, we took a mean of all temperature values collected during the baseline (last 30 s of the baseline, i.e. pre-test phase) by subject and session and then subtracted this value from all the temperature measurements taken during the post-test phase (30 s after stimulus offset). Finally, Mann-Whitney U tests were used to assess whether the temperature change for each individual from pre-test to post-test phase was significant. The resulting temperature change values could thus be statistically significantly negative, indicating a drop in nasal temperature between the pre-test and the post-test phase, and hence an increase in arousal as a result of the treatment. In this case, ‘increased arousal’ (coded as ‘1’) was attributed to the subject in the corresponding test and post-test phases. By contrast, a statistically significantly positive temperature change value indicated a rise in nasal temperature between the pre-test and post-test phase, and hence a decrease in arousal as a result of the treatment. In this case, ‘decreased arousal’ (coded as ‘-1’) was attributed to the subject in the corresponding test and post-test phases. Non-significant changes were attributed to ‘unchanged arousal’ (coded as ‘0’). Temperature data were available for 12–16 of the subjects for each treatment. Audio recordings were extracted from every exposure of the individuals to the treatments, while the temperature change was extracted only from one day due to time constraints in [35], resulting in missing data for arousal on some days were calls were recorded (coded as ‘NA’) (see further details including the procedure and results of Mann-Whitney U tests in [35]).

### Data collection

Vocalisations were recorded during each session at a distance of approximately 1 m from the vocalizing animal using a Sennheiser ME-67 directional microphone (frequency response: 40–20 000 Hz *±*2.5 dB) connected to a Marantz PMD-661 numeric recorder (sampling rate: 48 kHz; Kanagawa, Japan).

### Acoustic analysis

The analyses were restricted to calls produced during the test and post-test phases, and all vocalisations produced during these phases were extracted in Avisoft-SASLab Pro ver. 5.3.01. We then used Praat v.6.0.33 DSP Package [51] as well as Raven Pro 1.6 [52] and R version 4.2.3 [53] for parameter extraction, visualisation and further analyses.

For the purpose of the analyses, calls were manually classified as either a tsik, food call, Sd-peep, Sa-peep, P-peep, twitter, trillphee, phee, trill or ek based on their shape on the spectrogram (see [13] and [32] for descriptions). We excluded calls that overlapped with interfering background noise (e.g., calls of other animals, cage rattling, etc.) as well as short vocalisations for which the quality, after both visual examination of the spectrogram and listening to the audio recording, were insufficient to accurately determine whether the call was a tsik or a sd-peep call type.

We extracted both Mel-Frequency Cepstral Coefficients (MFCC) [54], as well as basic parameters from all vocalisations, resulting in 194 variables used in the analysis. MFCC (a total of 177 values) were extracted over nine time windows to account for differences in call duration (frequency range 5–10 kHz and 0–13 kHz for phees and eks, respectively, to reduce the effect of background noise) using R package BehaviouR [55]. Basic parameters included robust measurements obtained from Raven Pro (spectrogram settings: Hann window size = 10.7 ms, time grid overlap = 50%): Time 5%, Time 25%, Time 75%, Center time, Duration 50%, Duration 90%, Center frequency, Frequency 5%, Frequency 25%, Frequency 75%, Frequency 95%, Bandwidth 50%, Bandwidth 90%; which use the energy stored within the sound rather than boundaries of a selection making them less susceptible to user-related variations (for definitions see [56]). In addition, we extracted Amplitude Modulation (‘AM’) variation (cumulative amplitude variation per time unit), AM rate (number of complete cycles of amplitude modulation per time unit), AM extent (mean peak-to-peak variation of each amplitude modulation) [57], harmonicity, and mean Wiener entropy [58] in Praat. Extractions in Praat were made using a custom-built program, which batch processed the vocalisations, analysed the parameters and exported the data (adapted from [57–60]).

### Statistical analysis

The analyses were restricted to phees and eks, which were in sufficient number for analyses. All statistical analyses were carried out in R software on the two call types, phee and eks, separately, due to their highly distinct acoustic structure and function [13].

#### Support vector machines

We tested the ability of Support Vector Machines (SVM) to classify phee and ek calls according to valence (negative, neutral or positive), arousal level (−1 (significant increased arousal), 0 (no significant change), or 1 (significant decreased arousal)), food context (Yes/No) and social context (Yes/No). All measurements (basic and MFCC) were combined in this analysis. We only included calls that had a sample size equal to or greater than 15 per individual per class level. We then balanced these data (to account for potential individual differences) by randomly selecting 15 calls per individual per class category. We used a radial basis kernel type, with 5-fold cross validation with a modified version of the trainSVM function (GibbonR package [61]). The modification split the data 70:30 into a training and testing set without overlap and made it balanced per individual identity. Gamma and cost parameters were tuned automatically.

We ran separate models for each call type and separate models for valence, arousal, food context and playback. Accuracy was calculated from the confusion matrices (from models ran on the test (30%) set) with the function confusionMatrix (caret package [62]). As the calls selected for analysis were randomised and our sample sizes are not large, to avoid bias in results, we ran each model 5000 times. The reported accuracies are the mean values of the accuracies obtained from individual models. To assess whether the distribution of results was significantly different from random, we followed a onesided statistical approach. Specifically, we extracted the 5th percentile of the accuracy distribution for each model. If this value fell below the predefined random threshold (based on the number of classes), the model’s performance was considered not significantly different from random. Conversely, if all values above this threshold exceeded the random baseline, the result was deemed significant. This approach allowed us to determine whether classification accuracy was meaningfully better than chance.

#### Linear mixed effects models

A Principal Component Analysis (PCA) was first used to reduce redundancy among our vocal parameters, which were likely to be inter-related. To this aim, all parameters were z-transformed, and we then ran four PCAs in total (prcomp function, stats package), after ensuring suitability using a Bartlett’s sphericity test, on the MFCC and on the basic vocal parameters separately, and for each of the two call types. The scores of the first Principal Components (PC1) of each PCA were then extracted and used as response variables in Linear Mixed effects Models (LMMs) fit with Gaussian family distribution and identity link function (lmer function, lme4 package [63]).

In total, we carried out four LMMs, one for each call type and for each type of vocal parameter (MFCC and basic). Each LMM included the corresponding PC1 scores as the response variable, and four fixed effects (arousal, valence, food context and social context) without interactions. As LMMs can handle continuous factors, to test for the effect of arousal, we here (unlike for the SVM) included the actual average within-individual temperature change from pre-test to posttest phase (including all values, even non-significant changes; negative values indicated an arousal increase, and positive values an arousal decrease). Valence was entered as a three-level categorical fixed factor (negative, neutral or positive), while food and social contexts were entered as two-level categorical fixed factors (Yes/No) (Table 1). We included the identity and sex of the subjects, as well as the date of the test as a crossed random intercepts, to control for repeated measures of the same individuals and differences between sexes and days.

We checked for multicollinearity between our four fixed factors using the Variance Inflation Factor (‘VIF’; vif function, package car [64]). All values were within satisfactory range [65]; for arousal (temperature change; continuous factor), we verified that the generalised VIF (‘GVIF’) value was below 5–10, while for the categorical factors (valence, food context and playback), we verified that the value of GVIF^(1*/*(2*·Df*))^ was below SQRT(5 to 10), so below 2.24–3.16 [65, 66]. Arousal had a GVIF ranging from 1.15 to 1.30 depending on the models, while for the categorical factors, GVIF^(1*/*(2*·Df*))^ ranged from 1.91 to 2.95, except for the factor ‘food context’ that had a value of 3.31 for one model (including ek basic parameters as a response variable). Considering the statistical advantage of including all factors in the same model, we decided to keep all factors in despite this slightly high VIF value. The residuals of the LMMs were checked graphically for normal distribution and homoscedasticity (simulateResiduals function, package DHARMa [67]), and p values were calculated using a Type II Wald chi-square tests (Anova function, package car [64]). We extracted model estimate using the emmeans function (emmeans package [68]). If the valence (three-level categorical factor) had a significant effect, a Tukey HSD test (posthoc test) was used for further pairwise comparisons using the glht function (car package [64]). Finally, we extracted the marginal R^2^ (R2GLMM(m)) and conditional R^2^ (R2GLMM(c)) of our models using the r.squaredGLMM function (MuMIn package [69]). These two values were extracted both for the full models, as well as for significant factors by including the significant factor and random effects only. We considered effects significant if p *≤* 0.05, and marginally significant if 0.05 *<* p *≤* 0.06.

## Results

Each experimental session was divided into a baseline (“pretest”), a stimulation (“test”), and a post-stimulation (“posttest”) phase. During the pre- and post-test phases, the experimenter offered a small amount of food to the subject (mealworm or a small piece of nut cookie). The test phase differed between five treatments as follows (Table 1); 1) Control treatment (hereafter ‘Control’): no stimulus; 2) Food teasing (hereafter ‘Teasing’): the experimenter presented a highly appetitive food item (defrosted cricket (Acheta domesticus) or a nut cookie; i.e. an item of higher value than the food given during the pre- and post-test phases) to the subject, and pulled it back when the animal tried to grab it; 3) Agonistic calls (hereafter ‘PB-’): playback of agonistic calls (i.e. rapid fire tsik calls, tsik-ekk calls, alarm calls, twitter calls) produced by conspecifics from another group during intergroup visual contact; 4) Preferred food (hereafter ‘Food’): the subject was allowed to consume a defrosted cricket; 5) Food calls (hereafter ‘PB+’): playback of food calls [32, 38] (i.e. call ‘A’, ‘B’ and ‘C’ described in [32]) recorded from members of the subject’s social group while they were consuming preferred food. During the test phase of the two playback treatments and the Control, the experimenter turned their back to the subject and did not move. The change in emotional arousal of the subjects was assessed based on variation in nasal temperature measurements relative to baseline, while the putative valence was determined based on intuitive inference [39], following [35] (Table 1). The emotional valence (negative, neutral, positive) was assumed from the treatments based on intuitive inference [39] and knowledge of the species [35]; negative valence was assigned to the Teasing and PB-treatment, positive valence to the Food and PB+ treatment, and neutral valence to the Control treatment (Table 1).

We extracted all calls produced by the focal individuals during the test and post phases, but only phees and eks were emitted in sufficient numbers in all treatments to be included in our statistical analyses (Table 2). In total we extracted 625 phees (range = 3 to 153, mean = 44.64 *±* 37.23 calls per individual) and 594 eks (range = 1 to 174, mean = 45.69 *±* 46.17 calls per individual). Overall, 694 calls were extracted during the negative treatments (307 phees and 387 eks), 224 calls during the neutral treatments (Control) (127 phees and 97 eks), and 301 calls during the positive treatments (191 phees and 110 eks) (Table 2). From these extracted calls, we measured both Mel-Frequency Cepstral Coefficients (MFCC) and basic parameters (including frequency, duration and amplitude modulation parameters). MFCCs reflect the spectral envelope, which is mostly shaped by the vocal tract, while our basic parameters included parameters related to both the vocal folds (e.g. frequency parameters), vocal tract (e.g. energy quartiles and bandwidth), as well as time and amplitude domain (e.g. duration, amplitude modulation).

**Table 2.** Number of phee and ek calls included in our analyses as a function of the treatment, valence, food context, presence of sound played back and sex.

| Treatment | Sex | Call type |  |
| --- | --- | --- | --- |
|  |  | Phee | Ek |
| PB- | F | 52 | 79 |
| PB- | M | 99 | 191 |
| Teasing | F | 62 | 55 |
| Teasing | M | 94 | 62 |
| Control | F | 62 | 43 |
| Control | M | 65 | 54 |
| PB+ | F | 28 | 8 |
| PB+ | M | 48 | 61 |
| Food | F | 43 | 22 |
| Food | M | 72 | 19 |

### Support vector machines

First, we investigated the potential for the whole call structure (MFCC and basic parameters combined) to encode information about internal states (arousal change and valence), as well as external events (whether the context was food-related and whether it involved conspecific call playbacks (social context)), using support vector machines (SVMs). All SVMs classified both phees and eks to the considered factors (arousal change, valence, food context or social context) with accuracies exceeding chance levels (Table 3, Figure S1). Classifications based on valence were more accurate than classification based on arousal change, with mean accuracies of 64% vs. 57% for phees and 81% vs. 66% for eks, respectively. For phees, classification based on social context yielded higher accuracy (96%) than classification based on food context (75%). In contrast, for eks, classification accuracy was higher when distinguishing between treatments related to food or not (80%) than when distinguishing based on whether the treatment was social or not, i.e. involved conspecific call playbacks or not (68%).

**Table 3.**
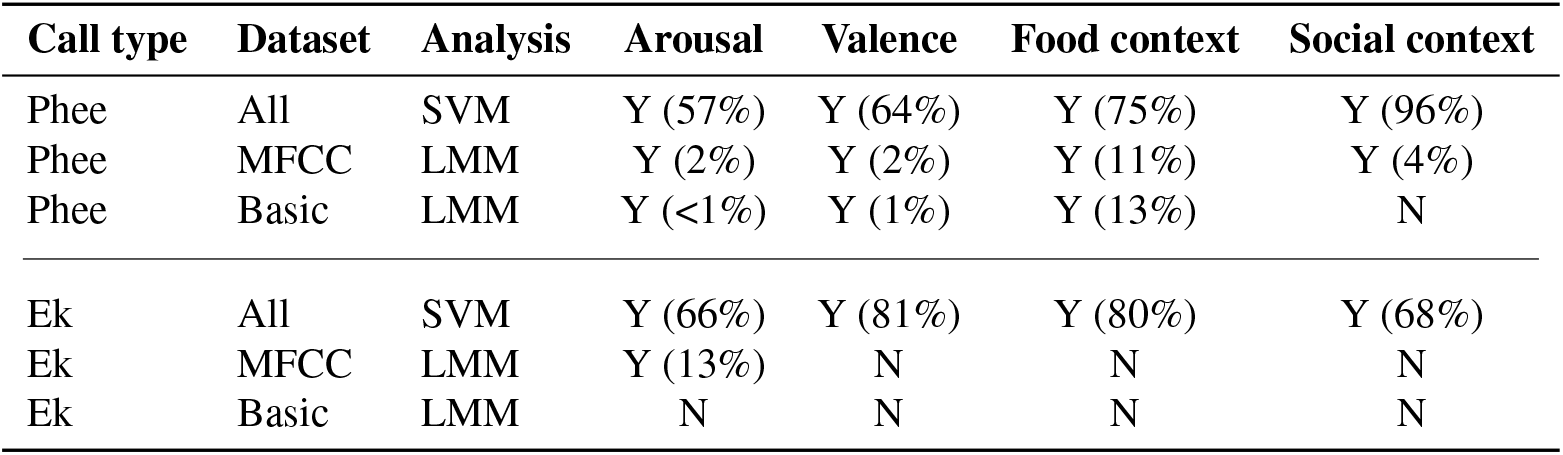
Overview of the results for phee and ek calls, for Support Vector Machines (SVM) and Linear Mixed Effects Models (LMM). Y = significant (i.e., higher than expected by chance for SVM), Y = marginally significant, N = not significant. The percentages indicated for significant results refer to the accuracy for SVM and effect sizes R2GLMM(m) for the LMM. Note that internal states are three-level classes (33% accuracy expected by chance for the SVM), while external factors are two-level classes (50% accuracy expected by chance in the SVM). SVM accuracy results are thus only comparable within call types between the two internal states, as well as between the two external factors.

| Call type | Dataset | Analysis | Arousal | Valence | Food context | Social context |
| --- | --- | --- | --- | --- | --- | --- |
| Phee | All | SVM | Y (57%) | Y (64%) | Y (75%) | Y (96%) |
| Phee | MFCC | LMM | Y (2%) | Y (2%) | Y (11%) | Y (4%) |
| Phee | Basic | LMM | Y (<1%) | Y (1%) | Y (13%) | N |
| Ek | All | SVM | Y (66%) | Y (81%) | Y (80%) | Y (68%) |
| Ek | MFCC | LMM | Y (13%) | N | N | N |
| Ek | Basic | LMM | N | N | N | N |

SVMs can be prone to spurious correlations and other hidden biases [40]. To corroborate these results, and further understand the effects revealed by the SVM, we used Linear Mixed effects Models (LMMs) to test whether MFCCs or basic parameters still encoded food context and/or social context, even after controlling for variation linked to arousal and valence. These models, unlike the SVMs, allowed us to test all factors in combination (i.e. in the same model) and to control for repeated measures as well as confounding factors such as the sex of the individuals. The LMM were carried out on the scores of the first principal component (PC1) extracted from a Principal Component Analysis (PCA) used, on each call type (phees and eks) and type of parameter (MFCC or basic parameters) separately, to reduce redundancy among our vocal parameters, which were likely to be inter-related.

#### Linear mixed effects models. Phee – MFCC

The PC1 extracted from the PCA carried out on phee MFCC parameters (‘PheePC1-MCFF’) explained 19.38% of the variance in the data (Table S1). Its values were significantly affected by all fixed factors: the arousal change (LMM: *χ*^2^ = 11.65, df = 1, p = 0.0006), the valence (χ^2^ = 9.38, df = 2, p = 0.009), whether the context was related to food or not (χ^2^ = 27.68, df = 1, p < 0.0001), as well as whether the context was social or not (χ^2^ = 12.49, df = 1, p = 0.0004) (full model: R2GLMM(m) = 10.16%, R2GLMM(c) = 75.98%; Figure 1A). PheePC1-MCFF decreased when temperature change increased (i.e. when arousal decreased) (R2GLMM(m) = 1.99%, R2GLMM(c) = 66.57%; Figure 1A). This effect remained significant after removing one outlier with a temperature change below -3 (Figure 1) (χ^2^ = 12.02, df = 1, p = 0.005). Concerning the effect of valence, post-hoc tests revealed that PheePC1-MFCC scores differed between the neutral treatment and both the negative and positive treatments (Tukey HSD test: p 0.025 for both), while they did not differ between the positive and negative treatments (Tukey HSD test: z = 2.21, p = 0.061) (R2GLMM(m) = 2.04%, R2GLMM(c) = 54.46%). Model estimates revealed that PheePC1-MFCC scores were overall significantly lower in the neutral treatment compared to both the negative treatments and positive treatments (Figure 1A). Treatments related to food obtained lower PheePC1-MCFF scores than those unrelated to food (R2GLMM(m) = 10.72%, R2GLMM(c) = 58.95%; Figure 1A), and, based on model estimates, treatments including playbacks (social contexts) obtained overall lower PheePC1-MCFF scores than other treatments (R2GLMM(m) = 3.62%, R2GLMM(c) = 53.92%; Figure 1A).

**Fig. 1.**
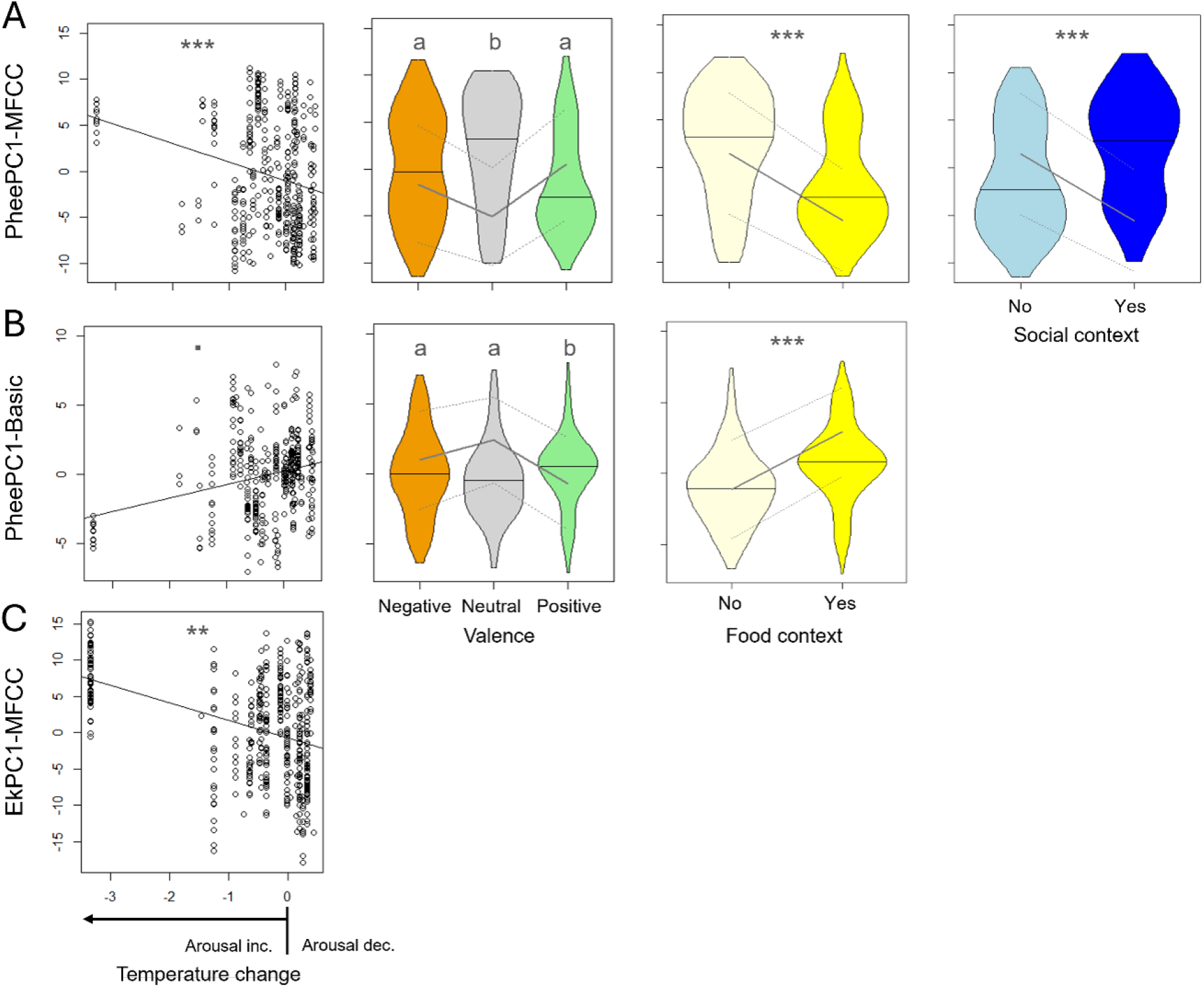
Results of the linear mixed effects models on Phee and Ek MFCC and Basic parameters. Only significant or marginally significant results are displayed: A) PheePC1-MCFF as a function of temperature change (arousal change: *<* 0 = arousal increase; 0 = no change; *>* 0 = arousal decrease), valence (negative (orange), neutral (grey) or positive (green)), food context (no (light yellow) = not related to food; yes (dark yellow) = related to food) and social context (no (light blue) = not social (no calls played back); yes (dark blue) = social (conspecific calls played back)); B) PheePC1-Basic as a function of arousal change, valence and food; C) EkPC1-MCFF as a function of arousal change. Plots: each point corresponds to a call; the black line indicates the regression line. Violin plot: the horizontal line shows the median, the grey lines show the model estimates (continuous line) and 95% confidence intervals (dashed lines). For each boxplot, different letters (a, b; Tukey HSD test) or stars (** 0.001 *≤* p *<* 0.01; *** *<* 0.001; linear mixed effects models) indicate significant differences (p *≤* 0.05), while ‘.’ Indicates a marginally significant result (0.05 *<* p *≤* 0.06). See also Figure 2 for PheePC1-MCFF and PheePC1-Basic as a function of the treatment.

### Phee – Basic parameters

The PC1 extracted from the PCA carried out on phee basic parameters (‘PheePC1-Basic’) explained 46.50% of the variance in the data (Table S2). This PC was negatively correlated with most vocal parameters (time of peak, duration, frequency quartiles, AM extent and Harmonicity), except for bandwidth parameters, AM rate and Wiener entropy, to which it was positively correlated (Table S2). Therefore, more positive (higher) PheePC1-Basic scores indicated sounds of lower frequencies, shorter durations, less tonal (more noisy), reaching peak frequencies earlier, and with wider bandwidths, and more but lower amplitude modulations. These scores were marginally significantly affected by arousal change (χ^2^ = 3.69, df = 1, p = 0.055), significantly affected by valence (χ^2^ = 12.51, df = 2, p = 0.002), as well as by whether the context was related to food or not (χ^2^ = 32.87, df = 1, p < 0.0001), but not by whether the context was social or not (χ^2^ = 2.46, df = 1, p = 0.12) (full model: R2GLMM(m) = 10.04%, R2GLMM(c) = 72.20%; Figure 1B). PheePC1-Basic scores tended to increase when temperature change increased (i.e. when arousal decreased) (Figure 1B), suggesting notably less tonal calls with lower frequencies and shorter durations when arousal decreased, although the R2 explained by temperature change alone was very low (R2GLMM(m) = 0.02%, R2GLMM(c) = 64.94%). This effect was, however, not marginally significant any longer after removing the outlier with a temperature change below - 3 (Figure 1) (χ^2^ = 2.83, df = 1, p = 0.093). Post-hoc tests for the effect of valence revealed that PheePC1-Basic scores differed between the positive treatment and both the negative treatments and neutral treatments (Tukey HSD test: p0.003 for both), while they did not differ between the negative and neutral treatments (Tukey HSD test: z = 1.94, p = 0.11) (R2GLMM(m) = 1.27%, R2GLMM(c) = 45.41%). Model estimates extracted from the LMM including all fixed and random factors showed that PheePC1-Basic scores were overall lower in the positive treatments, reflecting calls with notably higher frequencies and longer durations, compared to both the negative treatments and neutral treatments (Figure 1B). Finally, treatments related to food obtained higher PheePC1-Basic scores, suggesting calls with lower frequencies and shorter durations, than those produced in contexts unrelated to food (R2GLMM(m) = 12.50%, R2GLMM(c) = 55.15%; Figure 1B).

### Ek – MFCC

The first Principal Component (PC1) extracted from the PCA carried out on ek MFCC parameters (‘EkPC1-MCFF’) explained 28.31% of the variance in the data (Table S3). EkPC1-MCFF scores were significantly affected by the arousal change (LMM: χ^2^ = 8.92, df = 1, p = 0.003). However, they were not affected by the valence (χ^2^ = 0.94, df = 2, p = 0.62), by whether the context was related to food or not (χ^2^ = 1.14, df = 1, p = 0.29), nor by whether the context was social or not (χ^2^ = 0.0006, df = 1, p = 0.98) (full model: R2GLMM(m) = 19.34%, R2GLMM(c) = 63.40%; Figure 1C). EkPC1-MCFF decreased when temperature change increased (i.e. when arousal decreased) (R2GLMM(m) = 13.45%, R2GLMM(c) = 58.64%; Figure 1C). However, this arousal change effect was no longer significant after removing the outlier with a temperature change below -3 (Figure 1) (χ^2^ = 0.08, df = 1, p = 0.78).

### Ek – Basic parameters

PC1 extracted from the PCA carried out on ek basic parameters (‘EkPC1-Basic’) explained 42.50% of the variance in the data (Table S2). It was negatively correlated with parameters related to times of peak frequencies, duration and AM parameters, and positively correlated with parameters related to frequency quartiles (Table S2). Therefore, more positive (higher) EkPC1-Basic scores indicated sounds of shorter durations, reaching peak frequencies earlier and with less AM (AM var and AM extent), but characterised by higher frequencies. These scores were not affected by any of the factors; the arousal change (LMM: χ^2^ = 0.13, df = 1, p = 0.73), the valence (χ^2^ = 0.95, df = 2, p = 0.62), whether the context was related to food or not (χ^2^ = 0.09, df = 1, p = 0.77), and whether the context was social or not (χ^2^ = 1.27, df = 1, p = 0.26) (full model: R2GLMM(m) = 1.67%, R2GLMM(c) = 28.27%).

## Discussion

For a long time, animal vocalisations have been assumed to predominantly inform about internal states. It has been suggested that even if calls apparently encode information about external events, this may simply be the result of the internal states triggered by these events. This view has been increasingly questioned (e.g. [36, 37]), and the need to simultaneously measure internal states during call production has been emphasised and is increasingly implemented [36]. We therefore investigated how internal states (arousal and valence) as well as external events (food and social context) are encoded in the acoustic structure of common marmoset vocalisations, focusing on two call types that are given across diverse contexts: phees and eks. Overall, we found that phee calls encoded external events even when controlling for the variation linked to internal states, while eks reflected predominantly arousal change. A stronger encoding of external events in long-distance phee calls than in eks aligns with our predictions: in eks, which are given in close proximity, information about external events is less needed because it is aligned with all other individuals in proximity. Our results also fall in line with the conclusions of recent studies on marmoset vocalisations (e.g. [20, 32]), showing that common marmoset vocalisations, and particularly the phee contact calls, vary with external factors. Importantly, our results reveal that this is the case not only after controlling for arousal as previously shown [33], but also after considering emotional valence: phees encoded similar information both when the animals where handed over a highly desired food item (positive valence) and also when they were teased with it (negative valence).

Phee calls were accurately classified to all levels within internal states and external factors by the SVM (Table 3). SVMs are powerful tools to classify calls by each factor but are prone to spurious correlations [40] and less suitable to compare the relative importance of the four factors. We therefore complemented our analyses with LMMs, on both MFCCs and basic parameters to evaluate the robustness of the findings. These LMMs corroborated the SVM findings, and moreover revealed that food context explained more variation in the phee calls than all other factors. Furthermore, food context appears to be encoded more robustly than social context in the phees, since an effect was evident both in the basic acoustic parameters and the MFCCs, whereas an effect of social context was captured only in the MFCCs. Most notably, our food-related contexts included very diverse situations: an emotionally negative situation where food was present (food teasing), an emotionally positive situation in which food was also present (offering preferred food), as well as a situation where food was absent and subjects were simply exposed to a playback of conspecific food calls. Despite this diversity, the marmoset calls emitted in these food-related contexts differed considerably from those given in non-food-related contexts (Figure 2).

**Fig. 2.**
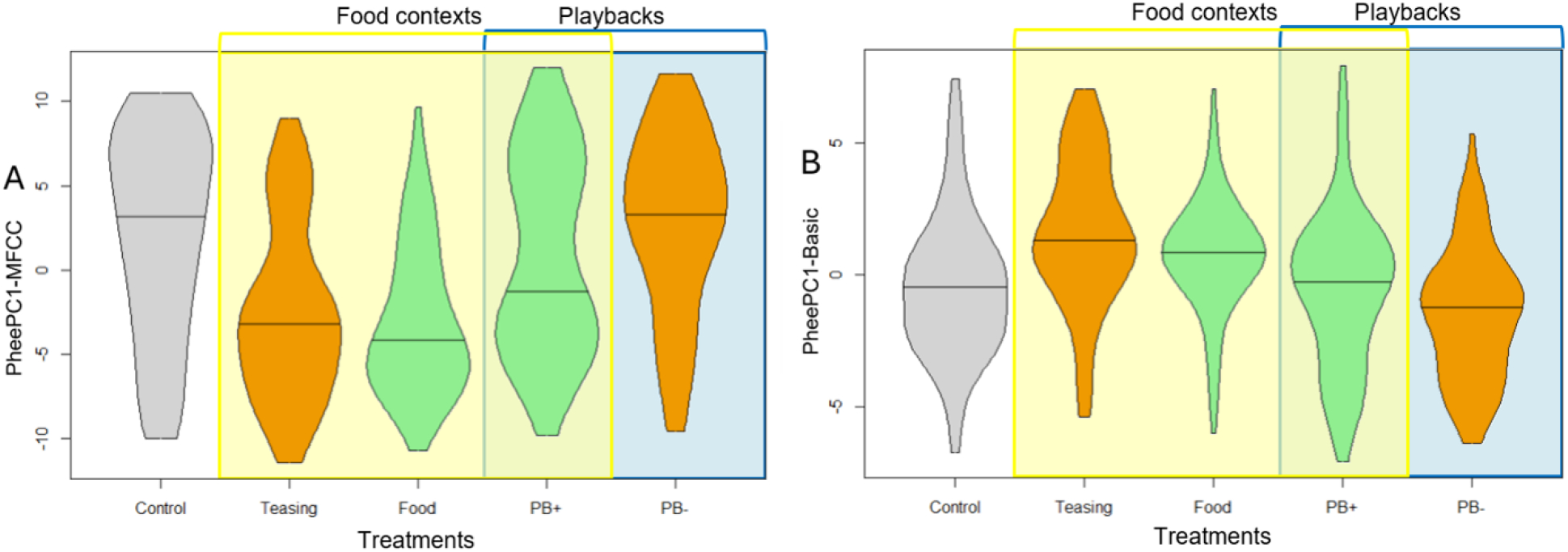
A) PheePC1-MCFF and B) PheePC1-Basic as a function of the treatment. Violin plot: the horizontal line shows the median. The colour code of the violin plots indicate the valence of the treatments (grey = neutral, orange = negative, green = positive). The food contexts (Teasing, Food and PB+) are highlighted in yellow and the social contexts (PB+ and PB-) in blue.

Eks (alert/mobbing calls) were also correctly classified by the SVMs at accuracy exceeding chance levels within all internal states and external factors. When analysing MFCC and basic parameters separately using LMMs, however, only arousal change significantly explained some variance in the MFCCs. Arousal might thus be a better predictor of the acoustic structure of ek calls than valence or external factors. Yet, this effect disappeared after removing one outlier. This suggests that arousal may play an even less pronounced role in shaping the acoustic structure of marmoset ek calls than our SVM and first LMMs showed. Future studies could test if arousal is not encoded in the rate of production of eks (i.e. number of eks per minute) rather than their acoustic structure.

Concerning emotion expressions, our analyses reveal both commonalities and differences with other species in how emotions are expressed in marmoset vocalisations. The SVMs, as well as the analysis of the basic parameters of phees calls, revealed effects of both arousal change (marginally significant) and valence. Notably, our results suggested that, with an increase in arousal, phees tended to become longer in duration, to be higher in frequency, more tonal, to peak in frequencies later, with narrower bandwidths, and less amplitude modulations that were smaller. An increase in duration and frequencies with arousal is commonly observed across species [2, 3]. However, across mammals, calls often become more chaotic (less tonal) when arousal increases, particularly in negatively valenced contexts [41]. Yet, this does not seem to apply to common marmosets, for which a negative correlation between heart rates and chaos (measured using entropy) was found across call types by Liao et al. [33]. Overall, the pattern we found aligns well with the results of this previous study [33], which found that phees produced between pairs of common marmoset became longer in duration, higher in dominant frequency, louder in amplitude and more tonal with increasing social distance and hence arousal. It should be noted, however, that the treatments we used only elicited very moderate amounts of arousal and negative valence, and calls emitted in more extreme situations are needed to complement our findings.

Regarding valence, based on our model estimates, positive valence also led to longer phees that were higher in frequency and more tonal, compared to neutral and negative valence. This is rather contradictory to the main trend found in other species, in which positive vocalisations tend to be shorter and with lower frequencies than negative ones [2, 3, 42]. One explanation for these disparities between our results and those found in other species could be that phee calls are not produced by vibration of the vocal folds, but by the apical vocal membranes [43]. Different modes of production might lead to different emotion-related changes in vocal parameters, as observed in encoding of arousal in rodent ultrasounds, which are emitted through laryngeal whistling [44], and often seem to deviate from the pattern observed across other vertebrates [45]. Further experiments testing how common marmoset express emotional valence in other call types and contexts will reveal if this is a common feature of this species.

Our results not only indicate that marmoset calls encode external events beyond the current emotional state of the emitter, but we also found that the way external events are parsed achieves an unexpected level of abstraction and context-independence: information about a food-related context seemed to be encoded in phee call regardless of whether the food was offered (positive situation), whether the animals were teased with food without getting it (negative situation), or whether they simply heard food calls from conspecifics (food absent) (Figure 2). This is both similar and different from human semantic communication, where arbitrary signs are used to refer to objects or events independently of the caller’s emotional state. While external contexts are encoded in calls, the marmosets are not using arbitrarily signs for this function. Instead, the call structure of phee calls changes (i.e. vocal accommodation). This falls well in line with a recent argument that vocal accommodation in fact paved the way toward vocal production learning in the narrow sense (Stoll et al. in rev.), because in contrast to the latter, across species, vocal accommodation is typically accompanied by several key features that characterise human language, such as babbling, extensive vocal turn-taking and high volubility. These key features are indeed highly prevalent in marmoset monkeys [26]. Playback experiments broadcasting phee calls recorded in the different food contexts will be essential to reveal if this information is perceived and relevant to others, and if others mentally represent food upon hearing these phee calls that encode food, similar to how this has been shown for food calls [32, 46].

## Conclusion

We found that common marmoset phee calls encode external events (food and social contexts) even when accounting for the variation linked to internal states (arousal and valence), while eks might better reflect arousal change. Interestingly, we showed that phee calls encode information about food in their call structure independently of their arousal (assessed through temperature changes), valence, and even food’s immediate presence. This might be akin, to some degree, to the phenomenon of ‘displacement’ described in human language (i.e. communication about events of objects that are not present in space or time [47, 48]). Marmoset phee calls not only encode the contexts studied here, but also the identity of the partner [20] as well as group membership, individual identity, and sex [15–17, 19]. Further work is not only needed to investigate whether marmoset listeners also successfully decode and use this information contained in phee calls, but also to fully disentangle how this rich information can be encoded simultaneously.

## Abbreviations

AM: Amplitude Modulation
MFCC: Mel-Frequency Cepstral Coefficients
SVM: Support Vector Machine
LMM: Linear Mixed effects Model
PCA: Principal Component Analysis
VIF: Variance Inflation Factor

## Declarations

### Ethics approval

The experiments described in the original publication [35], were approved by the Kantonales Veterinäramt des Kantons Zurich, Switzerland (license number 223/16).

### Consent for publication

Not applicable

### Availability of data and materials

All data and code used for the statistical analyses are available here https://zenodo.org/records/19729741.

### Competing interests

The authors declare that they have no competing interests.

### Funding

This project has received funding from the European Research Council (ERC) under the European Union’s Horizon 2020 research and innovation programme grant agreement No 101001295, the NCCR Evolving Language, Swiss National Science Foundation (SNF) Agreement no. 51NF40_180888, as well as Swiss National Science Foundation project funding 31003A_172979.

## Authors’ contributions

Study design: EFB, FAE, RKB and JMB; Data collection: FAE; Acoustic analyses: EFB, KW, EC and KM; Statistical analysis: EFB, KW, KSE; Supervision and funding: JMB; Writing – original draft: EFB and KW; Writing – review & editing: EFB, KW, JMB and RKB. All authors checked and approved the final version of the manuscript.

## Notes

https://zenodo.org/records/19729741

### Competing Interest Statement

The authors have declared no competing interest.

