## Supplementary Material for "What’s encoded in a marmoset phee call? Food context beyond arousal and valence"

**This PDF file includes:**

Fig. S1

Tables S1 to S3

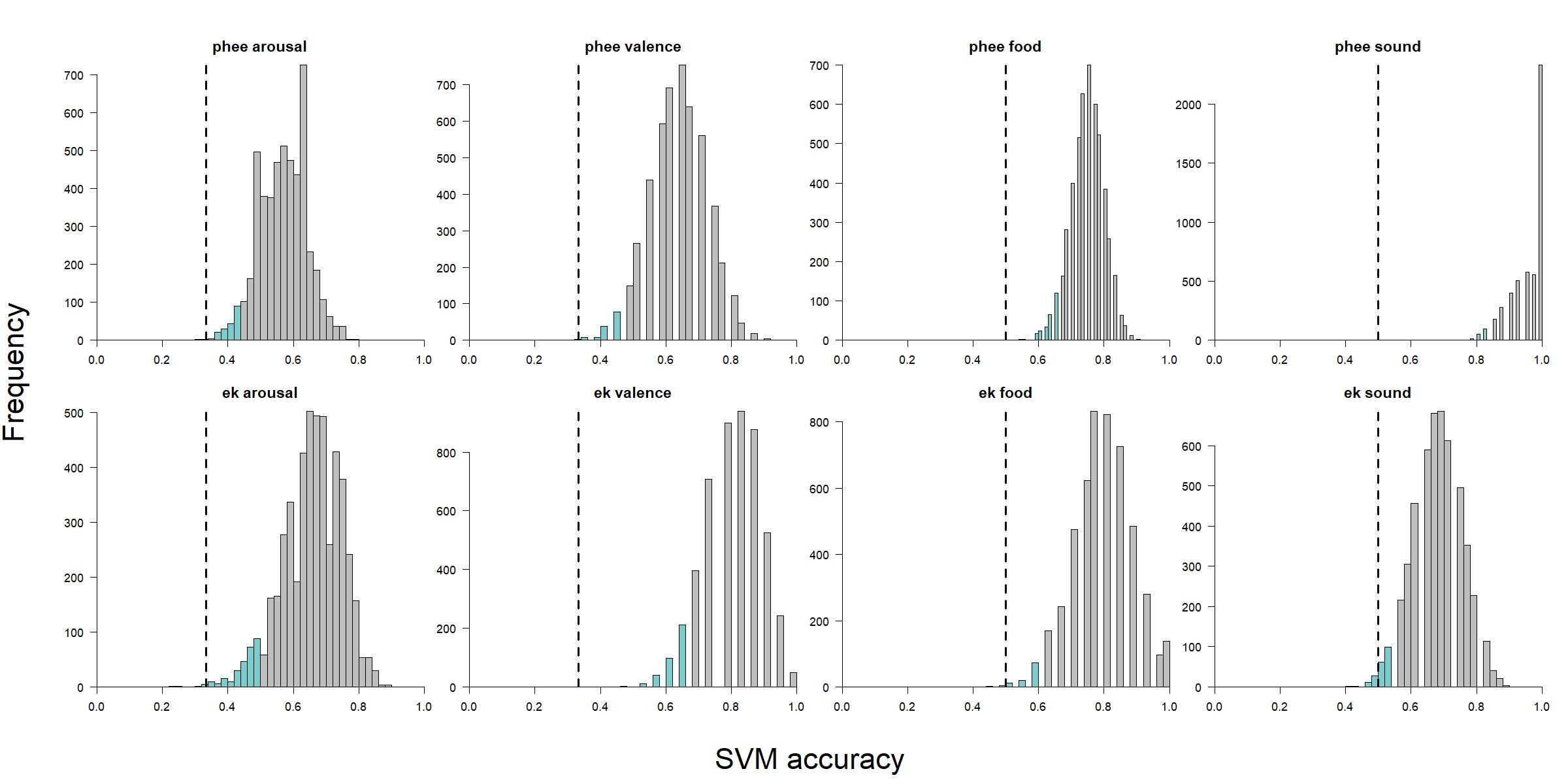

**Figure S1**. Distribution of SVM classification accuracy across models and conditions. Each panel shows the accuracy distribution for a classification task, with the bottom 5th percentile highlighted (blue). The dashed vertical line represents the predefined random threshold. If the 5th percentile falls below this threshold, the model's performance is not significantly different from random; otherwise, classification accuracy is considered better than chance.

**Table S1.** Loadings of the MFCC parameters (X1-X177) on the first principal component (PC1) of the principal component analysis carried out on Phee calls. Bold types indicate the heaviest factor loadings (|r| ≥ 0.50). See Table 1 for abbreviation of the parameters.

| **X1** | -0.46 | **X46** | 0.39 | **X91** | **-0.70** | **X136** | **-0.78** |
| --- | --- | --- | --- | --- | --- | --- | --- |
| **X2** | **-0.52** | **X47** | **0.68** | **X92** | **-0.53** | **X137** | 0.10 |
| **X3** | 0.46 | **X48** | **-0.53** | **X93** | 0.13 | **X138** | **-0.55** |
| **X4** | 0.42 | **X49** | -0.20 | **X94** | -0.17 | **X139** | -0.26 |
| **X5** | -0.12 | **X50** | 0.37 | **X95** | -0.48 | **X140** | 0.43 |
| **X6** | -0.31 | **X51** | -0.18 | **X96** | -0.14 | **X141** | 0.01 |
| **X7** | 0.02 | **X52** | -0.05 | **X97** | 0.18 | **X142** | -0.12 |
| **X8** | 0.13 | **X53** | 0.18 | **X98** | 0.13 | **X143** | 0.08 |
| **X9** | 0.00 | **X54** | -0.14 | **X99** | -0.05 | **X144** | **-0.83** |
| **X10** | 0.00 | **X55** | 0.00 | **X100** | **-0.75** | **X145** | **-0.77** |
| **X11** | -0.01 | **X56** | **-0.70** | **X101** | **-0.81** | **X146** | **-0.83** |
| **X12** | **-0.58** | **X57** | 0.48 | **X102** | **-0.74** | **X147** | **-0.73** |
| **X13** | -0.27 | **X58** | **0.56** | **X103** | **-0.66** | **X148** | 0.09 |
| **X14** | **0.66** | **X59** | **-0.62** | **X104** | -0.02 | **X149** | **-0.58** |
| **X15** | 0.27 | **X60** | 0.02 | **X105** | -0.34 | **X150** | -0.10 |
| **X16** | -0.45 | **X61** | 0.27 | **X106** | **-0.52** | **X151** | **0.50** |
| **X17** | -0.17 | **X62** | -0.24 | **X107** | 0.08 | **X152** | 0.02 |
| **X18** | 0.22 | **X63** | 0.08 | **X108** | 0.29 | **X153** | 0.03 |
| **X19** | 0.00 | **X64** | 0.09 | **X109** | 0.01 | **X154** | 0.18 |
| **X20** | -0.08 | **X65** | -0.15 | **X110** | -0.10 | **X155** | **-0.81** |
| **X21** | 0.04 | **X66** | 0.11 | **X111** | **-0.84** | **X156** | **-0.78** |
| **X22** | -0.01 | **X67** | **-0.72** | **X112** | **-0.81** | **X157** | **-0.83** |
| **X23** | **-0.65** | **X68** | **0.50** | **X113** | **-0.76** | **X158** | -0.68 |
| **X24** | 0.01 | **X69** | 0.47 | **X114** | **-0.75** | **X159** | 0.05 |
| **X25** | **0.75** | **X70** | **-0.60** | **X115** | -0.04 | **X160** | **-0.57** |
| **X26** | -0.07 | **X71** | 0.11 | **X116** | -0.39 | **X161** | -0.02 |
| **X27** | **-0.56** | **X72** | 0.14 | **X117** | **-0.51** | **X162** | 0.49 |
| **X28** | 0.13 | **X73** | -0.21 | **X118** | 0.19 | **X163** | 0.03 |
| **X29** | 0.18 | **X74** | 0.17 | **X119** | 0.14 | **X164** | 0.10 |
| **X30** | -0.16 | **X75** | -0.03 | **X120** | -0.21 | **X165** | 0.18 |
| **X31** | 0.04 | **X76** | -0.11 | **X121** | -0.17 | **X166** | **-0.74** |
| **X32** | 0.09 | **X77** | 0.14 | **X122** | **-0.87** | **X167** | **-0.75** |
| **X33** | -0.14 | **X78** | **-0.69** | **X123** | **-0.79** | **X168** | **-0.79** |
| **X34** | **-0.68** | **X79** | 0.32 | **X124** | **-0.78** | **X169** | **-0.65** |
| **X35** | 0.23 | **X80** | 0.47 | **X125** | **-0.79** | **X170** | 0.07 |
| **X36** | **0.75** | **X81** | **-0.50** | **X126** | 0.04 | **X171** | -0.43 |
| **X37** | -0.34 | **X82** | 0.12 | **X127** | -0.48 | **X172** | -0.10 |
| **X38** | -0.43 | **X83** | 0.04 | **X128** | -0.44 | **X173** | 0.43 |
| **X39** | 0.32 | **X84** | -0.13 | **X129** | 0.30 | **X174** | 0.04 |
| **X40** | 0.01 | **X85** | 0.13 | **X130** | 0.02 | **X175** | 0.12 |
| **X41** | -0.19 | **X86** | -0.07 | **X131** | -0.25 | **X176** | 0.14 |
| **X42** | 0.18 | **X87** | -0.02 | **X132** | -0.09 | **X177** | **0.70** |
| **X43** | -0.04 | **X88** | 0.13 | **X133** | **-0.86** |  |  |
| **X44** | -0.13 | **X89** | -0.57 | **X134** | **-0.77** | **Eigenvalue** | **5.86** |
| **X45** | **-0.69** | **X90** | **-0.70** | **X135** | **-0.82** | **% variance** | **19.38** |

**Table S2.** Loadings of the basic parameters on the first principal component (PC1) of the principal component analysis carried out on Phee and Ek calls. Bold types indicate the heaviest factor loadings (|r| ≥ 0.50). See Table 1 for abbreviation of the parameters.

|  | **Phee** | **Ek** |
| --- | --- | --- |
| Time.25...s. | **-0.72** | **-0.74** |
| Time.75...s. | **-0.79** | **-0.79** |
| Center.Time..s. | **-0.76** | **-0.82** |
| Dur.90...s. | **-0.75** | **-0.56** |
| Dur.50...s. | **-0.69** | **-0.70** |
| Center.Freq..Hz. | **-0.62** | **0.85** |
| Freq.5...Hz. | **-0.75** | **0.56** |
| Freq.25...Hz. | **-0.71** | **0.77** |
| Freq.75...Hz. | **-0.50** | **0.84** |
| Freq.95...Hz. | -0.33 | **0.76** |
| BW.50...Hz. | **0.50** | 0.47 |
| BW.90...Hz. | **0.67** | **0.66** |
| AM.var..dB.s. | 0.25 | **-0.77** |
| AM.rate..s.1. | **0.80** | -0.26 |
| AM.extent..dB. | **-0.66** | **-0.56** |
| Harmonicity | **-0.92** | -0.04 |
| mean.wiener.entropy | **0.82** | 0.19 |
| **Eigenvalue** | **2.81** | **2.69** |
| **% variance** | **46.50** | **42.50** |

**Table S3.** Loadings of the MFCC parameters (X1-X177) on the first principal component (PC1) of the principal component analysis carried out on Ek calls. Bold types indicate the heaviest factor loadings (|r| ≥ 0.50). See Table 1 for abbreviation of the parameters.

| **X1** | 0.00 | **X46** | -0.25 | **X91** | 0.13 | **X136** | 0.39 |
| --- | --- | --- | --- | --- | --- | --- | --- |
| **X2** | -0.22 | **X47** | 0.17 | **X92** | 0.36 | **X137** | 0.38 |
| **X3** | 0.26 | **X48** | -0.28 | **X93** | 0.25 | **X138** | -0.24 |
| **X4** | -0.22 | **X49** | 0.17 | **X94** | -0.30 | **X139** | **-0.82** |
| **X5** | 0.15 | **X50** | 0.41 | **X95** | **-0.80** | **X140** | **-0.89** |
| **X6** | 0.41 | **X51** | **0.69** | **X96** | **-0.83** | **X141** | **-0.92** |
| **X7** | **0.62** | **X52** | 0.31 | **X97** | **-0.86** | **X142** | **-0.90** |
| **X8** | 0.25 | **X53** | -0.14 | **X98** | **-0.82** | **X143** | **-0.85** |
| **X9** | -0.19 | **X54** | **-0.64** | **X99** | **-0.75** | **X144** | -0.40 |
| **X10** | **-0.62** | **X55** | **-0.85** | **X100** | -0.26 | **X145** | -0.27 |
| **X11** | **-0.76** | **X56** | -0.08 | **X101** | -0.14 | **X146** | 0.12 |
| **X12** | -0.06 | **X57** | -0.25 | **X102** | 0.20 | **X147** | 0.41 |
| **X13** | -0.24 | **X58** | 0.14 | **X103** | 0.46 | **X148** | 0.39 |
| **X14** | 0.24 | **X59** | -0.28 | **X104** | 0.36 | **X149** | -0.21 |
| **X15** | -0.24 | **X60** | 0.16 | **X105** | -0.33 | **X150** | **-0.79** |
| **X16** | 0.23 | **X61** | 0.44 | **X106** | **-0.84** | **X151** | **-0.88** |
| **X17** | 0.45 | **X62** | **0.69** | **X107** | **-0.88** | **X152** | **-0.91** |
| **X18** | **0.72** | **X63** | 0.33 | **X108** | **-0.91** | **X153** | **-0.90** |
| **X19** | 0.34 | **X64** | -0.15 | **X109** | **-0.88** | **X154** | **-0.84** |
| **X20** | -0.20 | **X65** | **-0.62** | **X110** | **-0.82** | **X155** | -0.38 |
| **X21** | **-0.67** | **X66** | **-0.84** | **X111** | -0.33 | **X156** | -0.25 |
| **X22** | **-0.83** | **X67** | -0.10 | **X112** | -0.21 | **X157** | 0.11 |
| **X23** | -0.07 | **X68** | -0.26 | **X113** | 0.12 | **X158** | 0.36 |
| **X24** | -0.25 | **X69** | 0.18 | **X114** | 0.38 | **X159** | 0.36 |
| **X25** | 0.24 | **X70** | -0.21 | **X115** | 0.34 | **X160** | -0.20 |
| **X26** | -0.29 | **X71** | 0.19 | **X116** | -0.31 | **X161** | **-0.78** |
| **X27** | 0.21 | **X72** | 0.41 | **X117** | **-0.83** | **X162** | **-0.86** |
| **X28** | 0.43 | **X73** | **0.65** | **X118** | **-0.89** | **X163** | **-0.90** |
| **X29** | **0.68** | **X74** | 0.29 | **X119** | **-0.92** | **X164** | **-0.89** |
| **X30** | 0.31 | **X75** | -0.11 | **X120** | **-0.89** | **X165** | **-0.83** |
| **X31** | -0.19 | **X76** | **-0.58** | **X121** | **-0.83** | **X166** | -0.39 |
| **X32** | **-0.66** | **X77** | **-0.84** | **X122** | -0.36 | **X167** | -0.28 |
| **X33** | **-0.85** | **X78** | -0.10 | **X123** | -0.24 | **X168** | 0.07 |
| **X34** | -0.08 | **X79** | -0.23 | **X124** | 0.13 | **X169** | 0.33 |
| **X35** | -0.24 | **X80** | 0.17 | **X125** | 0.40 | **X170** | 0.35 |
| **X36** | 0.20 | **X81** | -0.20 | **X126** | 0.38 | **X171** | -0.20 |
| **X37** | -0.29 | **X82** | 0.14 | **X127** | -0.28 | **X172** | **-0.78** |
| **X38** | 0.22 | **X83** | 0.38 | **X128** | **-0.84** | **X173** | **-0.84** |
| **X39** | 0.42 | **X84** | **0.64** | **X129** | **-0.90** | **X174** | **-0.89** |
| **X40** | **0.69** | **X85** | 0.28 | **X130** | **-0.92** | **X175** | **-0.87** |
| **X41** | 0.32 | **X86** | -0.10 | **X131** | **-0.90** | **X176** | **-0.82** |
| **X42** | -0.14 | **X87** | **-0.59** | **X132** | **-0.85** | **X177** | -0.04 |
| **X43** | **-0.67** | **X88** | **-0.82** | **X133** | -0.38 |  |  |
| **X44** | **-0.86** | **X89** | -0.22 | **X134** | -0.27 | **Eigenvalue** | **7.08** |
| **X45** | -0.08 | **X90** | -0.12 | **X135** | 0.09 | **% variance** | **28.31** |
